# MAFB surrogates the glucocorticoid receptor ability to induce tolerogenesis in dendritic cells

**DOI:** 10.1101/2021.07.27.453975

**Authors:** Octavio Morante-Palacios, Laura Ciudad, Raphael Micheroli, Carlos de la Calle-Fabregat, Tianlu Li, Gisela Barbisan, Miranda Houtman, Sam Edalat, Mojca Frank-Bertoncelj, Caroline Ospelt, Esteban Ballestar

**Affiliations:** Epigenetics and Immune Disease Group, Josep Carreras Research Institute (IJC), 08916 Badalona, Barcelona, Spain; Germans Trias i Pujol Research Institute (IGTP), 08916 Badalona, Barcelona, Spain; Center of Experimental Rheumatology, Department of Rheumatology, University Hospital Zurich, University of Zurich, Zurich, Switzerland.

**Keywords:** glucocorticoid receptor, MAFB, epigenetics, dendritic cells, tolerogenesis, inflammation

## Abstract

Glucocorticoids (GCs) exert potent anti-inflammatory effects in immune cells through the glucocorticoid receptor (GR). Dendritic cells (DCs), central actors for coordinating immune responses, acquire tolerogenic properties in response to GCs. Tolerogenic DCs (tolDCs) have emerged as a potential treatment for various inflammatory diseases. To date, the underlying cell type-specific regulatory mechanisms orchestrating GC-mediated acquisition of immunosuppressive properties remain poorly understood. In this study, we investigated the transcriptomic and epigenomic remodeling associated with differentiation to DCs in the presence of GCs. Our analysis demonstrates a major role of MAFB in this process, in synergy with GR. GR and MAFB both interact with methylcytosine dioxygenase TET2 and bind to genomic loci that undergo specific demethylation in tolDCs. We also show that the role of MAFB is more extensive, binding to thousands of genomic loci in tolDCs. Finally, MAFB knockdown erases the tolerogenic properties of tolDCs and reverts the specific DNA demethylation and gene upregulation. The preeminent role of MAFB is also demonstrated *in vivo* for myeloid cells from synovium in rheumatoid arthritis following GC treatment. Our results imply that, once directly activated by GR, MAFB takes over the main roles to orchestrate the epigenomic and transcriptomic remodeling that define the tolerogenic phenotype.

## Introduction

Dendritic cells (DCs) are a heterogeneous group of innate immune cells with a central role not only in the response to threats, but also in the regulation of inflammatory responses and the induction of immune tolerance. Several types of human DCs exist *in vivo*, including conventional DCs (cDCs) and plasmacytoid DCs (pDCs). Moreover, monocytes (MOs) can extravasate to tissues and differentiate to monocyte-derived macrophages (moMACs) or monocyte-derived dendritic cells (moDCs) ^1, 2^. MOs cultured with GM-CSF and IL-4 are a widely used *in vitro* model of DCs ^3^, with gene expression similarities with blood cDCs and *in vivo* moDCs ^4^.

Epigenetic determinants and transcription factors (TFs) critical to DC differentiation and function have been extensively studied. In particular, STAT6 ^5^, early growth response 2 (EGR2) ^6^, and interferon regulatory factor 4 (IRF4) ^4^ have been linked to DC fate determination and specific DNA demethylation events through recruitment of Ten-eleven translocation methylcytosine dioxygenase 2 (TET2), which is the most expressed TET enzyme in MOs ^7^. Moreover, aryl hydrocarbon receptor (AHR) can act as a molecular switch that enables monocyte differentiation to moDC, via IRF4, whereas MAFB determines monocyte differentiation to moMAC ^4, 8, 9^.

DCs can acquire tolerogenic functions *in vivo* and *in vitro* in response to several stimuli, including interleukin (IL)-10, vitamin D3, rapamycin, and glucocorticoids (GCs) ^10^. In particular, DCs differentiated from MOs *in vitro* with GM-CSF, IL-4, and GCs can suppress T cell proliferation *in vitro*, and display a high level of production of IL-10 and low levels of TNFα and IL-12p70 ^11, 12^. Tolerogenic DCs (tolDCs), generated as indicated above, can be useful as a treatment for autoimmune diseases. Several clinical trials have yielded satisfactory results in diseases such as rheumatoid arthritis ^13, 14^, multiple sclerosis ^15^, and Crohn’s disease ^16^.

GCs are a family of steroid hormones that are ligands of the Glucocorticoid Receptor (GR), a nuclear receptor expressed in most cell types that can trigger the expression of anti-inflammatory genes through direct DNA binding. Moreover, GR also represses the action of inflammatory-related TFs, such as the NF-KB and AP-1 families, via protein–protein interactions, in a process called transrepression ^17^. GR has been related to chromatin remodelers, including EP300 and BRG1 ^18, 19^. The mechanisms underlying cell type-specific programs induced by GR upon ligand binding, as well as the participation of TFs and epigenetic enzymes, remain to be fully determined, which is particularly relevant in the case of the various innate immune cell types.

In this work, we have studied the transcriptional and epigenetic remodeling associated with tolDC differentiation, and identified a major role for MAFB in this process, in synergy with GR. We have shown that GR binds both the promoter and enhancer regions associated with MAFB, which is quickly upregulated and binds thousands of genomic sites, correlating with widespread DNA demethylation and gene upregulation. We demonstrate how MAFB is crucial for the acquisition of the transcriptomic and epigenomic remodeling that gives rise to the tolerogenic phenotype. This is achieved through a differentiation switch to macrophage-like cells with tolerogenic properties. The major role of MAFB in activating a tolerogenic expression profile is also demonstrated in monocyte-derived cells from rheumatoid arthritis joints treated with GCs, in which an expansion of cells with a transcriptomic signature similar to MAFB-dependent tolDCs is shown.

## Results

### Dexamethasone modulates dendritic cell differentiation to a tolerogenic phenotype and drives transcriptome remodeling associated with MAFB

To investigate the mechanisms underlying the glucocorticoid (GC)-mediated phenotypic remodeling of dendritic cells (DCs), as well as the involvement of myeloid-specific TFs, monocytes (MOs) isolated from peripheral blood of healthy donors were differentiated *in vitro* to DCs and tolerogenic DCs (tolDCs) for 6 days using GM-CSF and IL-4 in the absence and presence of a GR ligand (dexamethasone), respectively (Figure 1A). CD8+ T cell proliferation assays in co-culture with both DCs and tolDCs revealed the immunosuppressive properties of the latter (Figure 1B). ELISA assays also showed that LPS+IFNγ-stimulated tolDCs produced higher levels of IL-10 and smaller amounts of TNFα, IL-12p70, and IL-1β than DCs (Figure 1C). Moreover, in the steady-state, higher IL-10 and lower IL-1β levels of production were observed in tolDCs, whereas TNFα and IL-12p70 were undetectable (Supplementary Figure 1A). tolDCs presented lower levels of expression of the costimulatory molecule CD86 and the maturation marker CD83 (Figure 1D). Moreover, tolDCs presented higher levels of CD14 and of the macrophage markers CD163 and CD16 (Supplementary Figure 1B).

**Figure 1.**
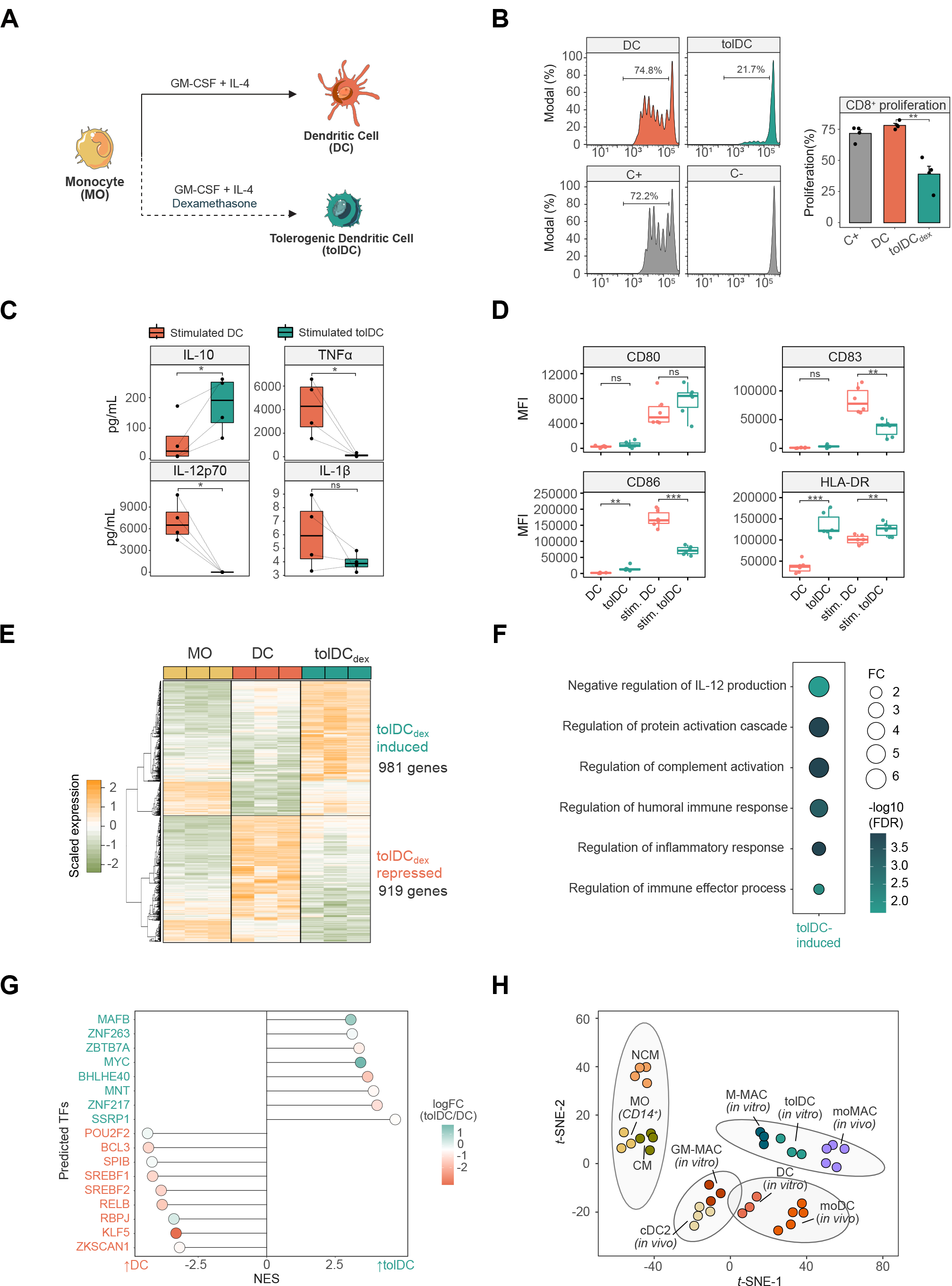
Phenotypic profiling of dexamethasone-mediated tolerogenic dendritic cells. (A) Schematic representation of the experimental approach, comparing dendritic cell (DC) with tolerogenic dendritic cell (tolDC) differentiation. (B) DC and tolDC were cocultured with CD8+ cells for 5 days. The final CFSE signal of CD8+ cells is shown (left panel). CD8+ with only CD3/CD28 T-activator beads (C+) or alone (C-) are also shown. In the right panel, the average proliferation of the quadruplicate is shown (mean ± standard error of the mean (SEM)). (C) IL-10, TNFα, IL-12p70 and IL1-β production of DC and tolDC, after 5 days of differentiation and 24 h of LPS (10 ng/μL) and IFNg (20 ng/μL) stimuli. P-values of paired t-tests are shown. (D) Box-plots of CD80, CD83, CD86 and HLA-DR surface expression (Median Fluorescence Intensity) in DCs and tolDCs in steady-state or stimulated with LPS (10 ng/μL) and IFNg (20 ng/μL) (ns p > 0.05, ** p < 0.01, *** p ≤ 0.001). (E) Gene expression heatmap of differentially expressed genes comparing tolDCs with DCs and also displaying the gene expression values of the precursor cell type (MO) (logFC > 0.5, FDR < 0.05). Scaled fluorescence values of expression arrays are shown, ranging from -2 (lower gene expression, green) to +2 (higher gene expression, orange). (F) Gene ontology (GO) over-representation of GO Biological Process categories. Fold change of tolDC induced genes over background and -log10(FDR) of Fisher’s exact tests are shown. (G) Discriminant regulon expression analysis (DoRothEA) of tolDC compared with DC. Only transcription factors with FDR < 0.05 are shown. NES and logFC of transcription factor expression are depicted. (H) T-distributed stochastic neighbor embedding (t-SNE) plot of the aggregated and batch-corrected gene expression data from our study (MO, DC and tolDC) and two additional public datasets (GSE40484 (moMAC, moDC, cDC2, CM (Classical MOs) and NCM (Non-Classical MOs) and GSE99056 (M-MAC (M2 Macrophages) and GM-MAC (M1 Macrophages)). The 4 different groups obtained using k-means clustering are represented with grey ellipses of multivariate t-distributions.

We then profiled the transcriptome of DCs, tolDCs, and MOs. 981 genes were induced and 919 genes were repressed in tolDCs in comparison with DCs (FDR < 0.05, logFC > 0.5) (Figure 1E and Supplementary Table 1). The transcriptomes of both tolDCs and DCs were notably dissimilar to that of MOs (Supplementary Figure 1C).

The Gene Ontology (GO) over-represented categories in tolDC-upregulated genes, including terms such as ‘negative regulation of IL-12 production’, ‘regulation of complement activation’, and ‘regulation of inflammatory response’ (Figure 1F). In tolDC-downregulated genes, terms such as ‘antigen processing and presentation of endogenous antigen’, ‘adaptive immune response’, ‘positive regulation of leukocyte activation’, and ‘response to interferon-gamma’ were over-represented (Supplementary Figure 1D).

tolDCs were enriched in several gene sets, including: ‘mo-MAC signature’ (genes upregulated in mo-MACs in comparison with mo-DCs) ^4^, genes upregulated in M-CSF macrophages relative to GM-CSF macrophages ^20^, and genes downregulated in response to GM-CSF and IL-4^21^ (Supplementary Figure 1E). On the other hand, tolDCs were depleted in inflammation-related gene sets such as Interferon-alpha response, TNFɑ Signaling via NF-κB genes upregulated in mo-DCs relative to mo-MACs)^4^, genes downregulated in M-CSF macrophages in comparison with GM-CSF macrophages, ^20^ and genes upregulated in response to GM-CSF and IL-4 cytokines ^21^ (Supplementary Figure 1F). We also confirmed the upregulation of canonical GR targets, such as PDK4, TSC22D3, ZBTB16, and FKBP5 in tolDCs, as expected (Supplementary Figure 1G).

To predict potential additional TFs involved in tolDC transcriptome acquisition, we performed a Discriminant Regulon Expression Analysis (DoRothEA). This analysis revealed that eight distinct TF regulons were enriched in tolDCs, although only two of these were associated with concomitant upregulation of their coding genes: *MAFB* and *MYC* (Figure 1G). To assess the possible role of MAFB in tolDC gene expression remodeling, we obtained a public dataset of genes downregulated after the treatment with an siRNA that targets *MAFB* ^8^ in M2 macrophages. These genes, positively regulated by MAFB, were more strongly expressed in tolDCs than in DCs, as can be observed in the GSEA (Supplementary Figure 1H). In addition, tolDCs presented a transcriptome similar to those of *in vitro* M2 macrophages and *in vivo* mo-MACs (Figure 1H). MAFB has previously been identified as a key mediator in the differentiation of both cell types ^4, 8^.

### Dexamethasone mediates the acquisition of a specific DNA methylation pattern which inversely correlates with gene expression during tolDC differentiation

In parallel, we obtained genome-wide DNA methylation profiles of MOs, DCs, and tolDCs after 6 days of differentiation. Principal component analysis (PCA) of all differentially methylated CpGs between groups showed non-overlapping clustering of MOs, DCs, and tolDCs alongside principal component 1 (Supplementary Figure 2A). Demethylation occurring during MO-to-DC differentiation was broadly inhibited in tolDCs (Supplementary Figure 2B).

The comparison of DCs and tolDCs (FDR < 0.05 and absolute Δß > 0.2) in relation to MOs revealed two clusters of CpG sites: a group of CpGs that underwent specific demethylation in DCs and that was blocked in tolDCs (C1, 1353 CpGs), and a second group that was specifically demethylated in tolDCs (C2, 411 CpGs) (Figure 2A and Supplementary Table 2).

**Figure 2.**
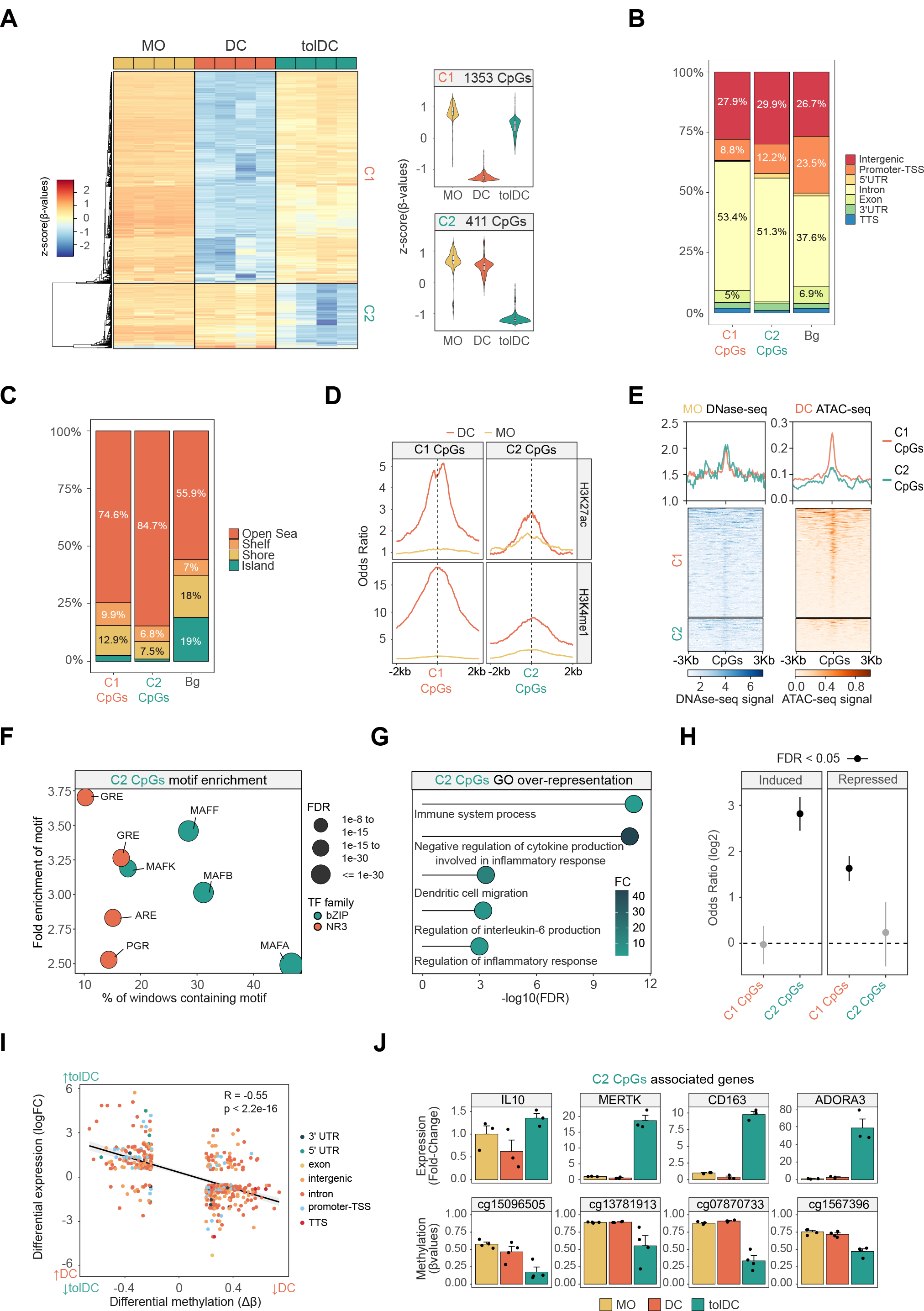
DNA methylation remodeling of dexamethasone-mediated tolerogenic dendritic cells. (A) DNA methylation heatmap of differentially methylated CpGs comparing DCs with tolDCs (Δβ ≥ 0.2, FDR < 0.05) and showing the DNA methylation values of the precursor cell type (MO). Scaled β-values are shown (lower and higher DNA methylation levels in blue and red, respectively). On the right side, violin plots of Cluster 1 (C1) and Cluster 2 (C2) depict scaled DNA methylation data. (B) Bar-plot of genomic features, percentages of C1 and C2 CpGs in comparison with background CpGs (Bg). (C) Bar-plot of CpG island contexts, percentages of C1 and C2 CpGs in comparison with background CpGs. (D) Accessibility (ATAC-seq) data of C1 CpGs (red) and C2 CpGs (blue) in MOs and DCs. The average DNAse-seq from an MO duplicate (BLUEprint) and ATAC-seq triplicate from DCs (GSE100374) were used in the representation. (E) ChIP-seq data of H3K27ac and H3K4me1 of CD14+ monocytes and DCs were downloaded from the BLUEPRINT database. Odds ratios of histone marks enrichment were calculated for bins of 10bp, 2000bp upstream and downstream in relation to C1 and C2 CpGs. CpGs annotated in the EPIC array were used as background. (F) Bubble scatter-plot of TF-binding motif enrichment for C2 CpGs. The x-axis shows the percentage of windows containing the motif and the y-axis shows the factor of enrichment of the motif over the EPIC background. Bubbles are colored according to the TF family. FDR is indicated by bubble size. (G) GO (Gene Ontology) over-represented categories in C2 CpGs. Fold change in comparison with background (EPIC array CpGs) and -log10(FDR) are shown. (H) C1 and C2 CpGs were associated with the nearest gene and the enrichment of both gene sets (C1 CpGs- and C2 CpGs-associated genes) over the tolDC-induced and tolDC-repressed genes were calculated using Fisher’s exact tests. Odds ratios ± 95% confidence intervals are shown. (I) DNA methylation of differentially methylated CpGs were correlated with gene expression of differentially expressed genes in the tolDC *vs.* DC comparison. LogFC of expression is represented in the y-axis, where a higher number represents a higher level of expression in tolDC and a lower number a higher level of expression in DC. DNA methylation is depicted on the x-axis as Δβ, where a lower number represents a lower level of methylation in tolDC, and a higher number a lower level of methylation in DC. Points are colored according to their genomic context. A significant negative correlation between methylation and expression is observed (R = -0.55, p < 2.2e-16). (J) The fold-changes of DC and tolDC expression (with respect to MO) of some examples of C2 CpGs associated genes are shown. Below each gene, the methylation (β-values) of an example of associated CpG is indicated.

Both CpG clusters were enriched in introns and depleted in promoters (Figure 2B). CpGs were generally located far from CpG islands (Open Sea) (Figure 2C). The two clusters were also enriched in enhancers and regions close to active transcription start sites (TSSs) (Supplementary Figure 2C). Looking at the enrichment in active enhancer histone marks (H3K4me1 and H3K27ac) in DCs and MOs, an increase in the signal of these marks was noted in MO-to-DC differentiation, especially in the C1, which became specifically demethylated in DCs (Figure 2D).

Employing the average signal of public MO DNAse-seq (Blueprint database)^22^ and DC ATAC-seq triplicates ^23^, we found that C1 and C2 CpGs had low accessibility in MOs. We also observed that C1 had greater accessibility than C2 in DCs, demonstrating an inverse correlation between chromatin accessibility and DNA demethylation (Figure 2E).

To detect potential TFs involved in the DNA methylation dynamics, we performed a TF motif enrichment analysis. As expected, C2 CpGs were enriched in Glucocorticoid Response Elements (GREs) and very closely related motifs (Androgen Receptor Elements, AREs, and Progesterone Receptor Elements, PGR). The Maf recognition element (MARE) was also enriched, including several TFs from the MAF family: MAFA, MAFB, MAFF and MAFK (Figure 2F). On the other hand, C1 CpGs were enriched in, among other binding motifs, AP-1 family, PU.1, IRF8, STAT6, and Egr1/Egr2 motifs (Supplementary Figure 2D), which are known to be associated with DC differentiation ^5, 6, 24, 25^.

Gene ontology (GO) analysis of C2 CpGs revealed enrichment of functional categories associated with immune system regulation, including ‘negative regulation of cytokine production involved in inflammatory response’ and ‘regulation of inflammatory response’ (Figure 2G). In contrast, GO analysis of C1 CpGs included categories related to inflammatory processes, such as ‘positive regulation of MAPK cascade’, ‘myeloid leukocyte activation’, and ‘inflammatory response’ (Supplementary Figure 2E). Moreover, C2 CpGs are demethylated in M2 macrophages (M-CSF), whereas C1 CpGs are demethylated in M1 macrophages (GM-CSF) (Supplementary Figure 2F).

We next associated each CpG with its nearest gene and tested whether C1 and C2 associated genes were enriched in tolDC-induced or tolDC-repressed genes. We found a strong enrichment of the induced genes over the C2 associated genes (FDR = 8.88e-41) and of the repressed genes over the C1 associated genes (FDR = 4.72e-26) (Figure 2H). In this regard, there was a significant inverse correlation between DNA methylation and gene expression (Figure 2I), which is exemplified in some CpGs associated with genes linked to the biology of tolDCs and DCs (Figure 2J, Supplementary Figure 2G).

In relation to the potential association between MAFB and our DNA methylation and expression data, we assessed the potential upregulation of MAFB-positively regulated genes of M2 macrophages and C2-associated genes in other tolDC types. In addition to tolDCs, we found that both gene sets were upregulated in DC10 (DCs differentiated *in vitro* from MOs with GM-CSF/IL-4 and IL10), but not in rapamycin-treated or vitamin D3-treated tolDCs (Supplementary Figure 2H).

### Dynamics of GR and MAFB genomic binding in tolDC differentiation

Since MAFB was predicted to be a TF of importance in the dexamethasone-specific gene expression (Figure 1H) and DNA methylation (Figure 2F) remodeling occurring in tolDCs, we investigated its role in the tolDC differentiation process and its interplay with GR.

*MAFB* expression was studied over time, where differences between DCs and tolDCs could be observed from 1 h of differentiation (Supplementary Figure 3A). MAFB protein levels could be detected by western blot in tolDC nuclei, at 12 and 24h, concomitant with GR translocation to the nucleus (Figure 3A). Moreover, by immunofluorescence at 24h of differentiation, MAFB was found to be localized in tolDC nuclei, whereas in DCs the level of expression was much lower (Figure 3B).

**Figure 3.**
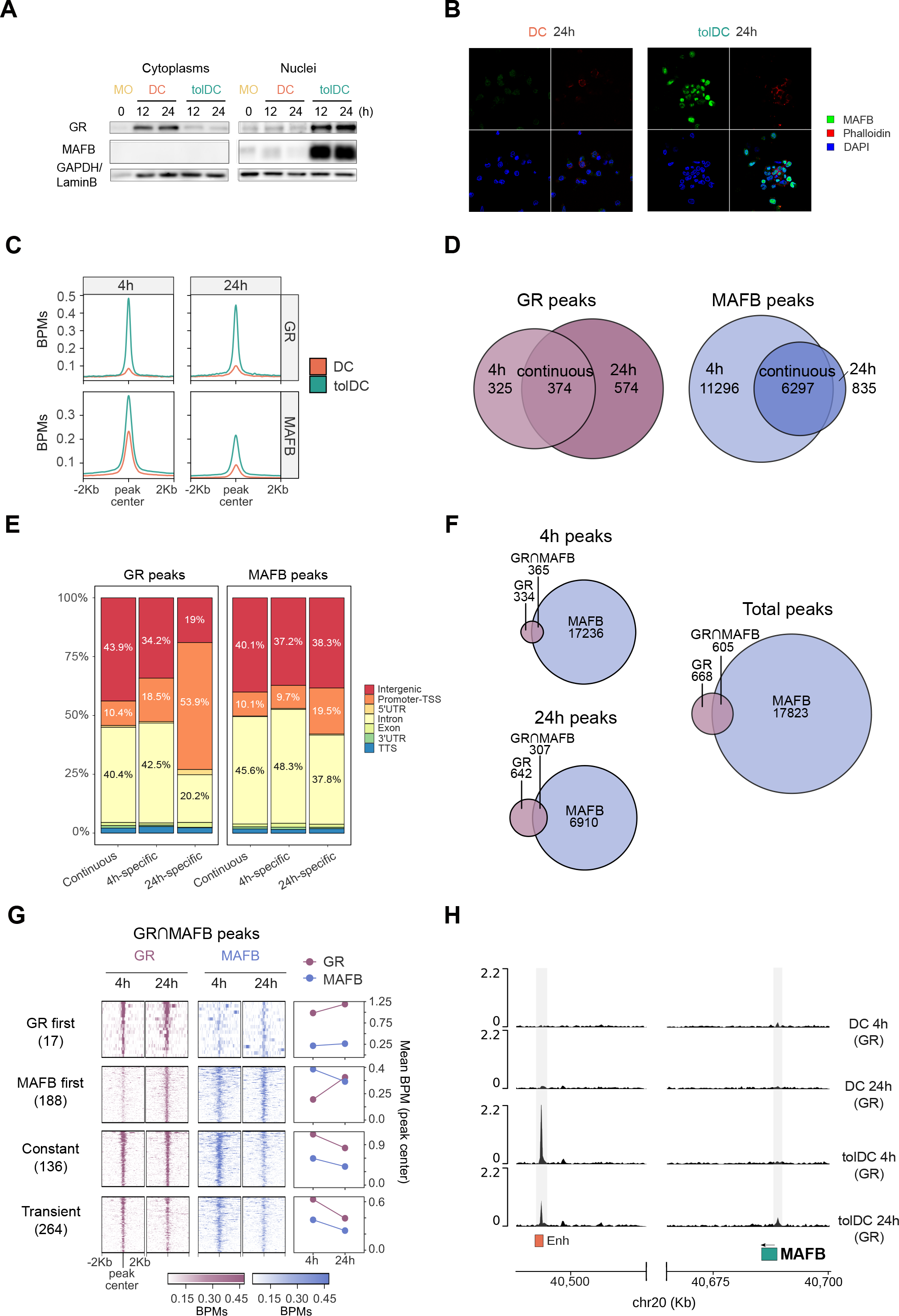
Delineation of GR and MAFB binding during DC and tolDC differentiation. (A) Western blot of glucocorticoid receptor (GR) and MAFB proteins in cytoplasm and nuclei of DC and tolDC. GAPDH and LaminB proteins were used as loading controls for cytoplasm and nuclei, respectively. (B) Immunofluorescence of MAFB in DC and tolDC after 24 h of differentiation. Fluorescent signal of MAFB (green), actin filaments (Phalloidin, red), nuclei (DAPI, blue) and a composite image with the sum of the three fluorescences are shown. (C) Average signal in bins per million mapped reads (BPMs) of GR and MAFB binding in DCs and tolDCs at 4 and 24 h of differentiation. (D) Venn diagrams showing GR and MAFB peaks at 4h, 24h and at both times (continuous). (E) Bar-plot of genomic features percentages of ‘4 h-specific’, ‘24 h-specific’, and ‘continuous’ peaks of GR and MAFB. (F) Venn diagrams showing GR and MAFB peaks at 4 and 24 h, and the concatenation of the two (total peaks). (G) Heatmap of total peaks with GR/MAFB overlap (GR⋂MAFB). Peaks were classified in ‘GR first’ (GR bound at 4 h and MAFB at 24 h), ‘MAFB first’ (MAFB bound at 4 h and GR at 24 h), ‘Constant’ (GR and MAFB peaks at 4 and 24 h), and ‘Transient’ (overlapping peaks not included in any of the former groups). (H) Representation of the GR ChIP-seq signal (BPMs) close to the MAFB gene. 15-states ChromHMM of MOs in peaks are depicted in orange. Significant peaks are shown against a grey background.

We next generated ChIP-seq data of GR and MAFB in DCs and tolDCs, at both 4h and 24h of differentiation. *De novo* motif discovery around GR and MAFB peaks in tolDC yielded very similar motifs to those of their respective canonical binding sites. (Supplementary Figure 3B). GR peak calling in tolDC resulted in hundreds of peaks with a strong signal (Figure 3C, Supplementary Figure 3C). By contrast, GR peak calling in DCs yielded a negligible number of significant peaks, at both 4h and 24h, with weaker signals (Figure 3C, Supplementary Figure 3C).

On the other hand, MAFB presented binding in thousands of sites in both DCs and tolDCs at 4h, with stronger signal and peak number in tolDCs (Figure 3C, Supplementary Figure 3C). Moreover, MAFB DC peaks at 4h were highly enriched in the MAFB canonical binding site (Supplementary Figure 3D) This is consistent with the preexisting expression of MAFB in MOs, which is downregulated during the MO to DC differentiation ^4^. In this regard, the MAFB peak number and signal in DCs at 24h is minimal in comparison with tolDCs, concordantly to a very low MAFB expression, and no motif compatible with the canonical one was found (Figure 3C, Supplementary Figure 3C, Supplementary Figure 3D).

To distinguish potential specific features of early and late peaks of GR and MAFB, we classified the tolDC peaks as ‘4h-specific’ (present at 4h but not at 24h), ‘24h-specific’ (present at 24h but not at 4h), or ‘continuous’ (present at both times) (Figure 3D and Supplementary Table 3). In both TFs, the subset with the strongest binding was the ‘continuous’ peaks. The binding typically occurred within open chromatin in MOs, even though some GR and MAFB peaks occurred in low-accessibility regions (Supplementary Figure 3E).

Whereas GR ‘4h-specific’ and ‘continuous’ peaks were located mainly in intergenic and intronic regions and were enriched in MO enhancers and active transcription start sites (TSS), ‘24h-specific’ peaks were most frequently found in promoters, and were more enriched in active TSS but not enriched in enhancers (Figure 3E, Supplementary Figure 3D). On the other hand, no notable differences were described in the location of MAFB ‘4h-specific’, ‘24h-specific’, and ‘continuous’ peaks, all of them being found mostly in intergenic and intronic regions, where they were enriched in monocyte active TSS and enhancers (Figure 3E, Supplementary Figure 3D).

Overall, there were many more MAFB than GR peaks, at 4h and 24h. Remarkably, GR binding, which was ubiquitously expressed in most cell types, significantly overlapped with the peaks observed in a panel of 53 GR ChIP-seqs (Supplementary Figure 3G), suggesting a common role of GR across these cell types.

We also inspected overlapping GR and MAFB peaks, which involved around half of all the GR peaks (Figure 3F). These GR∩MAFB peaks can be classified into four groups, depending on their temporal association between TFs: ‘GR first’, where GR peaks at 4h, and MAFB peaks at 24h, but not at 4h; ‘MAFB first’, where MAFB peaks at 4h, and GR peaks at 24h, but not at 4h; ‘constant’, where GR and MAFB peaks at both 4 and 24h; and ‘transient’, overlapping peaks not included in any of the former groups (Figure 3G). In general, ‘GR first’, ‘constant’ and ‘transient’ peaks were enriched in GREs. In contrast, ‘MAFB first’ peaks, in which MAFB is bound before GR, are enriched in ETS family motifs, including PU.1, and AP-1 motifs, but not MAREs. (Supplementary Figure 3H). All groups were enriched in MO enhancers and TSSs, except for ‘MAFB first’ peaks, which were not enriched in enhancers (Supplementary Figure 3I).

To address the mechanism of the GC-induced upregulation of MAFB, we checked the binding of GR around the MAFB promoter. At both 4h and 24h, GR was bound in an enhancer in MOs whose closest gene was MAFB. At 24h, GR was also bound to the MAFB gene promoter. (Figure 3G). This rapid binding, together with the rapid MAFB RNA and protein upregulation (Figure 3A, Supplementary Figure 3A), suggests a direct, GR-mediated, mechanism of MAFB regulation.

### GR and MAFB binding are correlated with gene upregulation and DNA demethylation in tolDCs

We measured the association between each ChIP-seq peak and its nearest gene in order to examine the link between TF binding and gene expression remodeling. Overall, both MAFB and GR ‘4h-specific’, ‘24h-specific’, and ‘continuous’ associated genes were more strongly expressed in tolDC than in DCs, indicating that both TFs may be involved in upregulating tolDC -specific genes (Figure 4A, Figure 4B). It is notable that, in both TFs, the most frequently associated subset was the ‘continuous’ associated genes.

**Figure 4.**
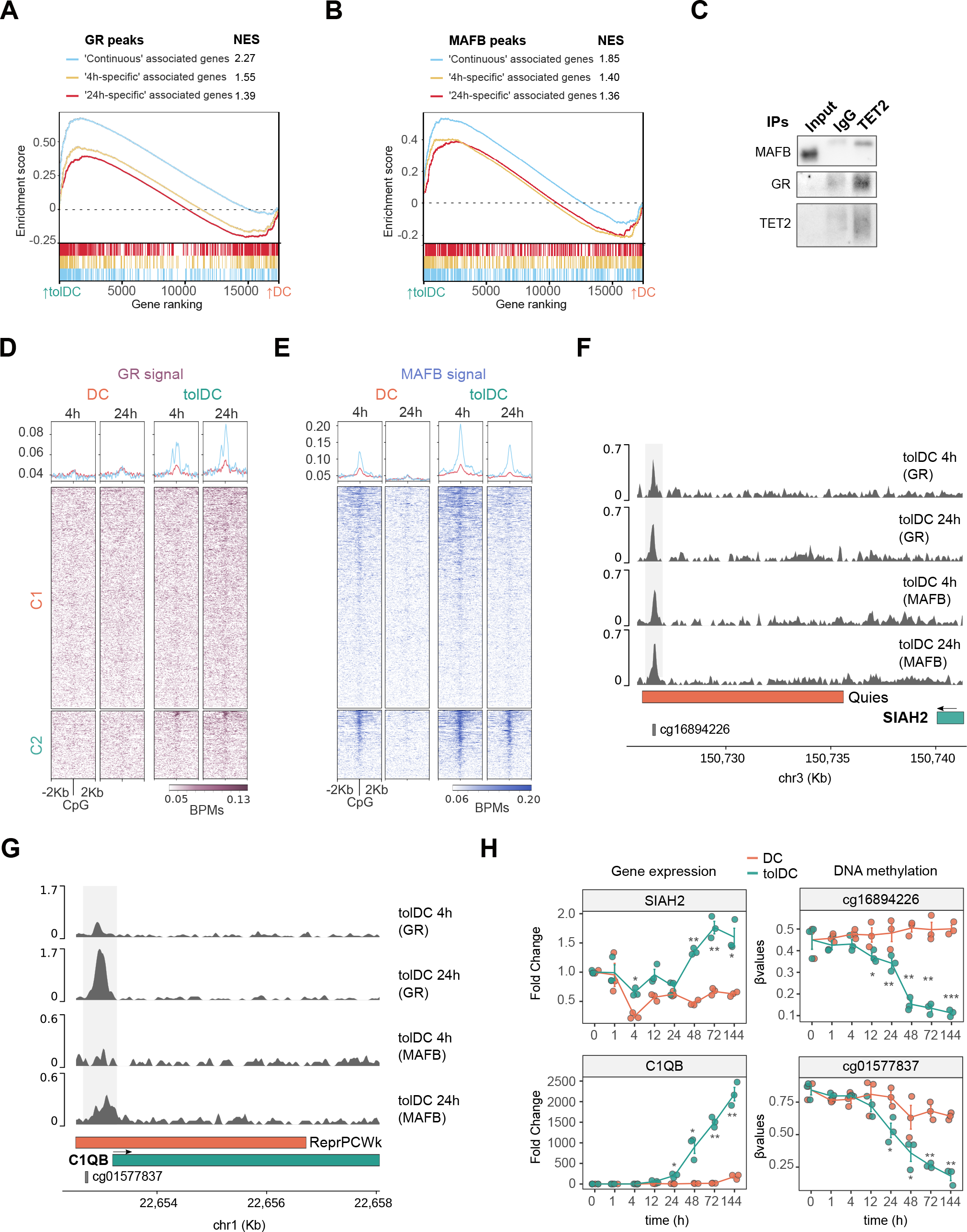
Integration of GR and MAFB binding with DNA methylation and gene expression changes in tolDCs. (A) Gene set enrichment analysis (GSEA) of tolDC *vs.* DC, with the genes closest to GR ‘continuous’, ‘4h-specific’ and ‘24h-specific’ peaks. The running enrichment score is presented with the normalized enrichment score (NES) is shown above (FDR < 0.01). (B) Gene set enrichment analysis (GSEA) of tolDC *vs.* DC, with the genes closest to the top500 MAFB ‘continuous’, ‘4h-specific’ and ‘24h-specific’ peaks (500 peaks with the highest joint p-value). The running enrichment score is represented in the y-axis and the normalized enrichment score (NES) is shown above (FDR < 0.01). (C) Western blot of the co-immunoprecipitation of TET2, showing the signal of MAFB, GR and TET2 proteins. (D-E) Heatmaps of the GR and MAFB ChIP-seq signal (BPMs) around C1- and C2-CpGs in DCs and tolDCs at 4 h and 24 h of differentiation. (F-G) Representation of the GR and MAFB ChIP-seq signal (BPMs) close to the SIAH2 and C1QB genes. 15-state ChromHMM of MOs in peaks are depicted in orange. Significant peaks are shown against a grey background. (H) SIAH2 and C1QB time-course gene expression (relative arbitrary units) and DNA methylation (β-values) of associated differentially methylated CpGs.

To determine the relationship between TF binding and DNA methylation, we first profiled the initial methylation state of CpGs surrounding GR and MAFB peaks, using public whole-genome bisulfite sequencing data of CD14+ MOs ^22^. Both TFs bound methylated regions, even though they were located mainly in non-methylated regions (Supplementary Figure 4A). However, GR 24h-specific peaks were found almost exclusively in non-methylated regions, concordantly with its enrichment in MO active TSS (Supplementary Figure 3F).

We then performed co-immunoprecipitation of TET2, a key mediator of active demethylation in myeloid cells, revealing its interaction with both GR and MAFB (Figure 4C) in tolDCs. In this regard, GR ChIP-seq signal in tolDCs at both 4h and 24h was found around some CpGs specifically demethylated in tolDCs (C2 CpGs), with a strong signal around a small subset of total C2 CpGs (Figure 4D). Moreover, MAFB signal in tolDCs at both 4h and 24h was found more generally around C2 CpGs. Interestingly, MAFB signal was also present in DC at 4h, at a lower level than that of tolDCs, but was not present at 24h (Figure 4E).

C1 CpGs were associated with EGR2, among other TFs (Supplementary Figure 2D), which prompted us to use a public dataset of EGR2-FLAG ChIP-seq of DCs ^6^, which revealed a strong signal in C1 CpGs but not in C2 CpGs (Supplementary Figure 4B). Strikingly, the EGR2 gene was downregulated in tolDCs (Supplementary Figure 2G), which may partially explain the blockage of C1 CpG demethylation. In addition, the genes closest to the EGR2 peaks were more strongly expressed in DCs, and downregulated in tolDCs, suggesting an association between a potential loss-of-function of EGR2 in tolDCs and the specific downregulation of genes (Supplementary Figure 4C).

We also studied the temporal relation between DNA methylation and gene expression changes. We selected CpGs with GR and/or MAFB binding associated with tolDC-induced or tolDC-repressed genes (Figure 4F, Figure 4G, Supplementary Figure 4D, Supplementary Figure 4E, Supplementary Figure 4F). In some loci, DNA demethylation was concomitant with or preceded gene upregulation (Figure 4H). However, there were also examples where gene upregulation clearly anticipated DNA demethylation (Supplementary Figure 4G).

### MAFB downregulation reverts dexamethasone-induced expression and DNA methylation remodeling, damping the tolerogenic phenotype of tolDCs

Given the association between MAFB binding, gene upregulation and DNA demethylation in tolDCs, we performed MAFB knockdown, using small-interfering RNAs (siRNAs) targeted against MAFB (siMAFB) or non-targeting (siCTL). Under our conditions, we achieved around 75% transfection efficiency (Supplementary Figure 5A), more than 50% RNA reduction (Supplementary Figure 5B) and a drastic decrease in MAFB protein (Supplementary Figure 5C).

Gene expression profiling of tolDCs transfected with siMAFB or siCTL was performed by RNA-seq, obtaining 222 downregulated genes and 259 upregulated genes (FDR < 0.05, logFC > 0.5) (Figure 5A and Supplementary Table 4). Among the downregulated genes, relevant tolDC-induced genes were found, including *RNASE1*, *CCL18*, *LGMN*, *MERTK*, *IL10*, *IFIT3* and *SLCO2B1*. Moreover, the upregulated genes included tolDC-repressed genes such as *GPT*, *CD1C*, *TNF*, *EGR2*, *CSF2RB*, *CD1B* and *FLT3*.

**Figure 5.**
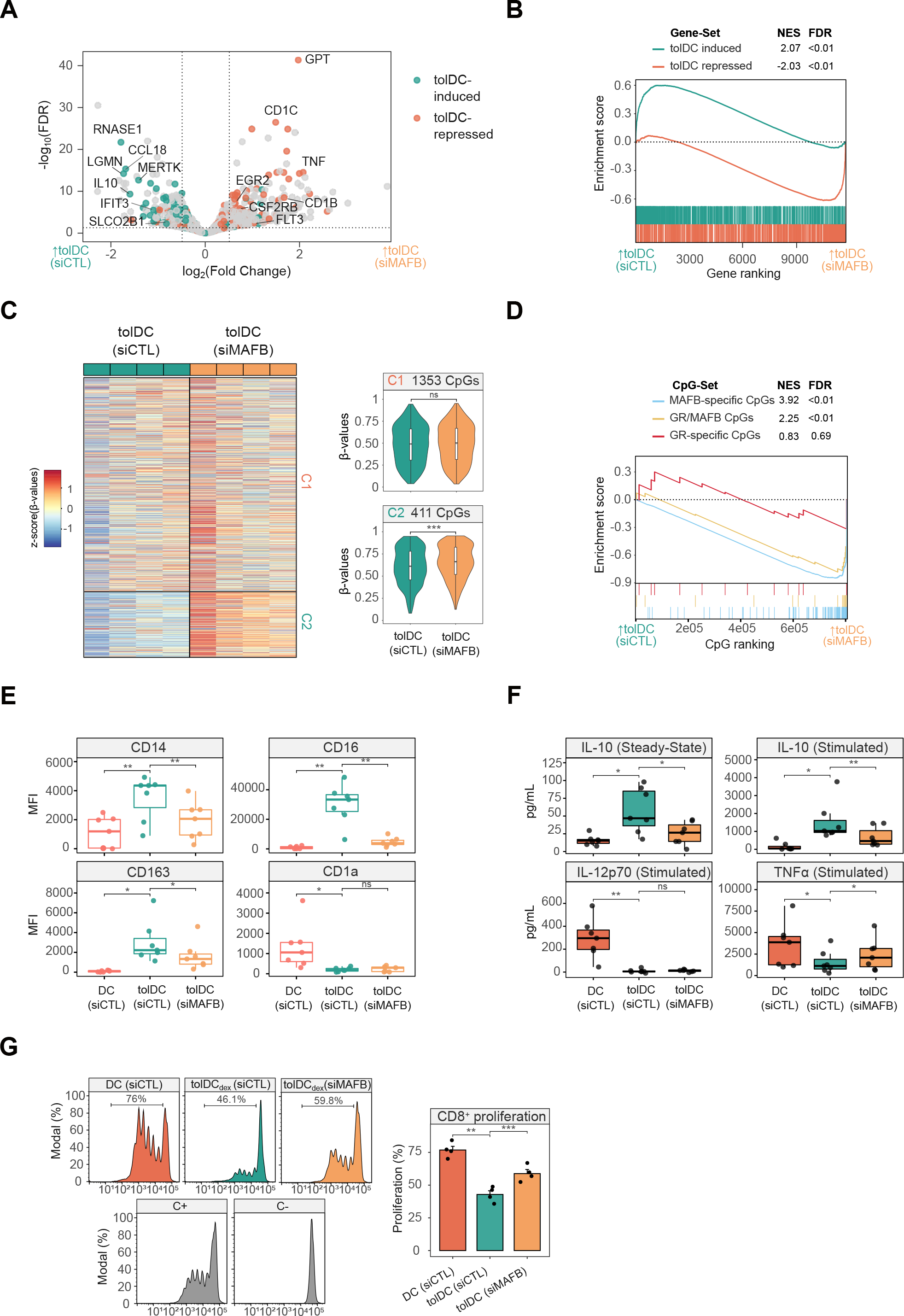
Effects of MAFB knockdown during tolDC differentiation. (A) Volcano plot comparing tolDCs treated with control siRNA (siCTL) and MAFB siRNA (siMAFB). Dashed lines indicate significance thresholds (FDR < 0.05, absolute logFC > 0.5). tolDC-induced and tolDC-repressed genes are shown in blue and orange, respectively. (B) Gene set enrichment analysis (GSEA) of tolDCs (siCTL) *vs.* tolDCs (siMAFB), using tolDC-induced and tolDC-repressed gene sets. The running enrichment score is represented and the normalized enrichment score (NES) is shown above (FDR < 0.01). (C) DNA methylation heatmap of previously obtained differentially methylated CpGs (C1-CpGs and C2-CpGs) in tolDCs (siCTL) and tolDCs (siMAFB). Scaled β-values are shown (lower DNA methylation levels in blue and higher methylation levels in red). On the right side, violin plots of Cluster 1 (C1) and Cluster 2 (C2) depict β-values (ns p > 0.05, *** p ≤ 0.001). (D) Methylated CpG set enrichment analysis (mCSEA) of tolDCs (siCTL) *vs.* tolDCs (siMAFB), using MAFB-only CpGs, GR/MAFB CpGs and GR-only CpGs as CpG-sets (depending on the overlap of CpGs with GR or MAFB peaks). The running enrichment score is represented and the normalized enrichment score (NES) and FDR are shown above. (E) Box-plots of median fluorescence intensity (MFI) of CD14, CD16, CD163 and CD1a flow cytometry data from DCs (siCTL), tolDCs (siCTL) and tolDCs (siMAFB) (n = 7) (ns p > 0.05, * p < 0.05, ** p ≤ 0.01). (F) Box-plots of supernatant concentration from DCs (siCTL), tolDCs (siCTL) and tolDCs (siMAFB) (n = 7) of IL-10 in steady-state and stimulated conditions (LPS 10 ng/μL and IFNγ 20 ng/μL) and IL-12p70 and TNFα under stimulated conditions (pg/mL). TNFα and IL-12p70 in steady state were not detected. (ns p > 0.05, * p < 0.05, ** p ≤ 0.01) (G) DC (siCTL), tolDC (siCTL) and tolDC (siMAFB) were cocultured with CD8+ cells for 5 days (n = 4). The final CFSE signal of CD8+ cells is shown (left panel). CD8+ with only CD3/CD28 T-activator beads (C+) or alone (C-) are also shown. On the right panel, the average proliferation of the quadruplicate is shown (mean ± standard error of the mean (SEM)) (** p ≤ 0.01, *** p ≤ 0.001).

GO Biological Process-over-represented categories in siMAFB-downregulated genes included terms such as ‘monocyte chemotaxis’, ‘regulation of tolerance induction’ and ‘negative regulation of interferon-gamma production’ (Supplementary Figure 5D). Furthermore, among siMAFB-upregulated genes, terms such as antigen processing and presentation via MHC class Ib, positive regulation of leukocyte-mediated immunity and leukocyte differentiation were over-represented (Supplementary Figure 5E).

In this regard, tolDC-induced genes were, in general, significantly downregulated with the MAFB siRNA, whereas tolDC-repressed genes were upregulated (Figure 5B). Moreover, genes associated with MAFB ‘continuous’ and ‘4h-specific’ ChIP-seq peaks were linked to siMAFB downregulation, whereas those GR peaks were not related to downregulation or upregulation (Supplementary Figure 5F).

We then tested the effect of MAFB downregulation on the differentially methylated CpGs. CpGs specifically demethylated in tolDCs (C2 CpGs) were more methylated in tolDC when MAFB was downregulated, confirming the role of MAFB in the tolDC demethylation process (Figure 5C). In contrast, no differences were observed in C1 CpGs, corresponding to the absence of MAREs in the cluster (Supplementary Figure 2B) and the weaker signal of MAFB in the ChIP-seq (Figure 4D). Based on their overlap with MAFB and GR ChIP-seq peaks, C2 CpGs were then divided into three groups: MAFB-specific, GR/MAFB, and GR-specific CpGs. In siMAFB-treated tolDCs, MAFB-specific and GR/MAFB CpGs were more methylated, whereas GR-specific CpGs were not affected by MAFB inhibition (Figure 5D). This suggests that both MAFB and GR can direct demethylation to C2 CpGs, consistent with their interaction with TET2.

Surface markers CD14, CD16, and CD163 were significantly reduced in tolDCs with the MAFB inhibition (Figure 5E), providing evidence of the functional role of MAFB in the tolerogenic phenotype.

Moreover, the inhibition of MAFB also reduced IL-10 production at steady-state and after stimulation of the cells with LPS. TNFα production in stimulated cells was significantly higher in siMAFB-treated tolDCs, showing that MAFB inhibition not only reduced the tolerogenic features of tolDCs but also boosted an increase in some proinflammatory traits, concordantly with RNA-seq data (Figure 5F).

Consequently, the suppression of CD8+ T cell proliferation, a main feature of tolDC, was reduced with the MAFB inhibition (Figure 5G). Overall, these data indicate that MAFB is a key player in the acquisition of tolDC tolerogenesis, and in the transcriptomic and epigenomic events driving that phenotype.

### Glucocorticoids skew an MAFB-associated, monocyte-derived cell differentiation program in rheumatoid arthritis joints

*In vitro* tolDCs and DCs derived from MOs resembled *in vivo* cell types, moMACs and moDCs, respectively (Figure 1H). We therefore performed single-cell RNA-seq from unsorted cells of the synovium of rheumatoid arthritis (RA) joints treated, or not, with GCs to further explore the *in vivo* effects of GCs in monocyte-derived cell populations.

From the total single-cell transcriptomes, we exclusively selected putative MOs and MO-derived cells, based on the expression of *CD14*, *S100A9*, and *MRC1* ^1^, and the absence of *THY1*, *CD248*, *CD27, IGHN*, *CD3G*, *CD3E*, *CD34*, *KLRD1* and *NKG7* expression (Supplementary Figure 6A-C). Clustering of these cells yielded five distinct subpopulations. We then excluded clusters containing fewer than 50 cells and annotated the three remaining clusters as M1, M2 and M3 (Figure 6A).

**Figure 6.**
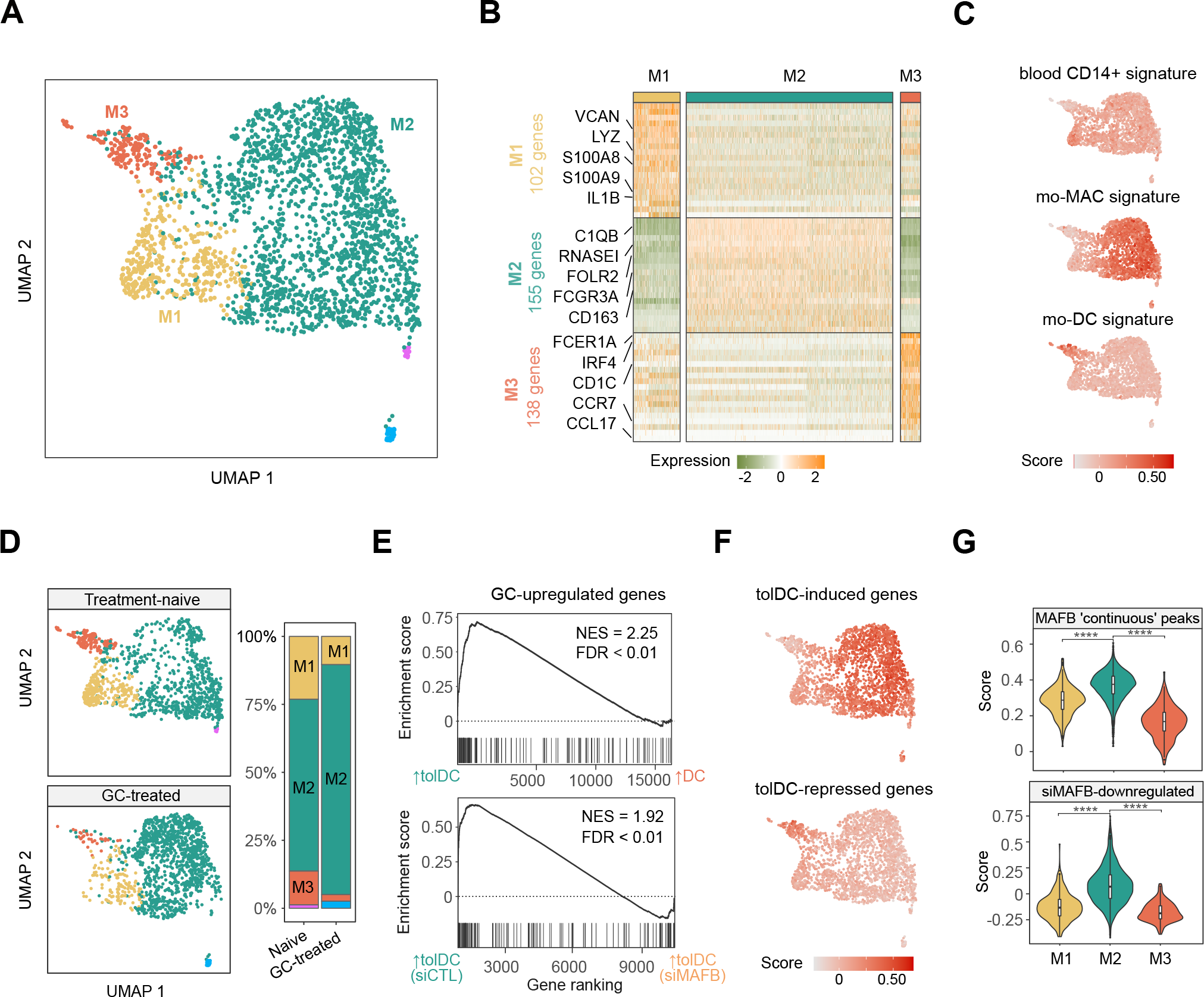
*In vivo* effects of glucocorticoid treatment in myeloid cells from RA synovium. (A) UMAP of putative monocyte-derived clusters from synovial tissues identified in the scRNA-seq analysis. (B) Heatmap of the top 20 genes most differentially expressed from the M1, M2 and M3 clusters. Relevant cluster markers and the total number of genes identified in each cluster are shown. (B) UMAP heatmap of module scores of gene sets associated with blood CD14⁺ cells, mo-MACs and mo-DCs ^65^. (D) UMAP is divided depending on the treatment of RA patients (treatment-naive or GC-treated). In the right panel, proportions of each cluster in each group are shown. (E) Gene set enrichment analysis (GSEA) of tolDCs *vs.* DCs and tolDCs (siCTL) *vs.* tolDCs (siMAFB), using the genes upregulated with glucocorticoids in M1, M2 and M3 cluster cells from patients as the gene set. The running enrichment score is represented and the normalized enrichment score (NES) is shown above (FDR < 0.01). (F) UMAP heatmap of module scores of tolDC-induced and tolDC-repressed genes. (G) Violin plots of module scores of genes associated with MAFB ‘continuous’ peaks and siMAFB-downregulated genes in the M1, M2 and M3 clusters (**** p ≤ 0.0001).

Looking at the gene markers with a high level of expression in each group, we found some monocytic and proinflammatory genes, such as VCAN, LYZ, S100A8, S100A9 and IL1B, in the M1 cluster, macrophagic markers (C1QB, FCGR3A, CD163, RNASE1 and FOLR2) in the M2 cluster, and dendritic cell-related markers (FCER1A, IRF4, CD1C, CCR7 and CCL17) in the M3 cluster (Figure 6B).

In addition, module scores of blood CD14⁺ MO, mo-MAC and mo-DC signatures displayed an increased signal in M1, M2 and M3 clusters, respectively (Figure 6C).

We then analyzed the relative proportions of clusters in the treatment-naive and the GC-treated patient joints. In the GC-treated patient joints, M1 and M3 clusters were depleted, whereas the M2 cluster was increased (Figure 6D). In this regard, when M1, M2 and M3 clusters expression were analyzed in bulk, GC-upregulated genes were more expressed in *in vitro* tolDCs and downregulated with the MAFB siRNA (Figure 6E).

Moreover, *in vitro* tolDC-induced genes presented a higher level of expression in the M2 cluster, whereas tolDC-repressed genes were more strongly expressed in M3 (Figure 6F). Furthermore, the module scores of the genes associated with MAFB ‘continuous’ ChIP-seq peaks and the siMAFB-downregulated genes for each cluster showed an increased signal in the M2 cluster for both modules (Figure 6G), supporting the involvement of MAFB in the transcriptomic signature of the M2 cluster.

## Discussion

In this study, we demonstrate a fundamental role for MAFB, in combination with GR, in the acquisition of glucocorticoid-mediated tolerogenesis by tolDCs. We show a coordinated action of MAFB and GR in their binding to genomic sites, DNA methylation and gene expression changes, which lead to the establishment of a tolerogenic phenotype, in which MAFB plays a predominant role. This major role for MAFB is confirmed *in vivo*, by examining single-cell data from the synovium of rheumatoid arthritis patients treated with GC.

GR, as a ubiquitously expressed nuclear factor, is distributed fundamentally in the cytoplasm of cells, until GCs induce a conformational change and promote its translocation to the nucleus and binding to the genome in a matter of minutes^26^. This, together with the previously described pioneer capacity of GR^27^, suggested a direct and major role in the acquisition of the tolDC phenotype and methylome. On the other hand, MAFB has been previously linked to MO differentiation ^9^ and the M2 macrophage phenotype ^8^. Strikingly, we show that GR has an important but limited direct role in this context, given that its binding occurs in a few hundred genomic loci and is associated with a small fraction of the epigenomic and transcriptomic remodeling produced. Instead, after the glucocorticoid stimulus, GR binds to an enhancer close to *MAFB*, and the upregulation of the gene is observed after as little as 1h of differentiation, pointing to a direct, GR-mediated regulation mechanism. MAFB, in turn, binds thousands of genomic loci in tolDCs, and is involved not only in DNA demethylation and gene upregulation, but also in the acquisition of the tolerogenic phenotype, as demonstrated by its knockdown. In this regard, MAFB acts as a surrogate to induce tolerogenesis in DCs on behalf of GR.

MAFB is a known downstream target of the IL-10/STAT3 signaling pathway^28^, and additional evidence of the major role of MAFB in our model comes from the finding that *in vitro* DC-10 (tolDCs generated with IL-10) present upregulation of MAFB target genes and C2 CpG-associated genes, despite the absence of active GR.

A central question about GR biology has been how a TF that is ubiquitously expressed across most tissues can have different cell-type-specific functions^29–31^. Chromatin accessibility has been described as pre-determining GR binding and shaping its differential binding across various cell types^32^. Our data also indicate a preference for GR to bind preexisting open chromatin. However, we found a significant overlap of tolDC GR peaks across very different cell lines. Our findings support that, upon GC exposure, MAFB confers the cell type specificity required to acquire the tolerogenic phenotype of DCs.

DNA methylation (5mC) is generally considered a repressive epigenetic mark that is associated with gene downregulation^33^. TET2, the most strongly expressed TET in MOs, has been linked to active demethylation events in these cells in several terminal myeloid differentiation models ^5–7, 34^. We prove that GR and MAFB both interact with TET2 in tolDCs, indicating that the demethylation process is probably triggered by MAFB- or GR-driven TET2 recruitment. However, we cannot rule out the possibility that other TET proteins are involved.

DNA methylation changes occurring during the MO-to-tolDC differentiation is potentially both a cause and a consequence of the reshaping of gene expression. Some examples show that DNA demethylation can precede gene upregulation, or can occur at the same time (SIAH2, C1QB), whereas other genes, such as CD163 and ETS2, present a noticeable gene upregulation before DNA demethylation, as shown in other immune contexts^35^. The functional role of the latter is enigmatic. Increasing evidence suggests that active demethylation intermediates, such as 5hmC, may be epigenetic marks with regulatory functions ^36^. DNA demethylation could also have a role later on, stabilizing the phenotype or fine-tuning the immune response after successive inflammatory stimuli. This is compatible with *IL10* gene behavior; whereby dexamethasone-mediated demethylation precedes a higher level of production of IL-10 after an inflammatory stimulus.

The transcriptome of GM-CSF/IL-4 DCs is very similar to the *in vivo* mo-DCs described in ascitic and synovial fluids ^4^. Intriguingly, tolDCs are more similar to *in vivo* moMACs and to *in vitro* M2 macrophages. MAFB upregulation is a hallmark of these three cell types. Several studies have produced evidence to suggest that there is a macrophagic phenotype of dexamethasone-treated monocytes, and that an increase of phagocytosis is a typical feature of tolDCs ^37, 38^. Here, we show that glucocorticoids skew MO-to-DC differentiation through MAFB, resembling an *in vivo* cell type. In this regard, we have shown depletion of cells with an expression pattern similar to moDCs and an increase of cells similar to moMACs in RA patients treated with GCs. Since *in vivo* moDCs are involved in the pathogenesis of several inflammatory diseases ^2, 39, 40^, the GC-mediated remodeling of monocyte-derived populations in the synovium could be a significant process that modifies their proportions and modulates the inflammation produced by monocyte-derived cells in tissues.

Our results shed light on the regulatory mechanisms of GC-induced tolDC differentiation, identifying the critical role of MAFB, which takes over GR to fulfil the main roles of tolerogenesis induction. We have described a new mechanism of action of GCs in MOs, consisting of MAFB induction, overriding the MO-to-DC differentiation program, rendering macrophage-like tolerogenic cells. Moreover, in MO-derived cells from synovial tissues, we have also shown a concordant GC-mediated depletion of mo-DCs and an increase of mo-MACs. By improving the understanding of the molecular mechanisms underlying the tolDC generation mediated by GCs and the effects of GCs in MOs *in vivo*, our results can provide insights to enhance the *in vitro* generation of tolDCs and to create more specific anti-inflammatory therapies.

## Methods

### CD14+ monocyte purification and culture

Buffy coats were obtained from anonymous donors via the Catalan Blood and Tissue Bank (CBTB). The CBTB follows the principles of the World Medical Association (WMA) Declaration of Helsinki. Before providing blood samples, all donors received detailed oral and written information and signed a consent form at the CBTB.

PBMCs were isolated by density-gradient centrifugation using lymphocyte-isolation solution (Rafer). Pure MOs were then isolated from PBMCs by positive selection with magnetic CD14 MicroBeads (Miltenyi Biotec). Purity was verified by flow cytometry, which yielded more than 90% of CD14⁺ cells.

MOs were resuspended in Roswell Park Memorial Institute (RPMI) Medium 1640 + GlutaMAX™ (Gibco, ThermoFisher) and immediately added to cell culture plates. After 20 minutes, monocytes were attached to the cell culture plates, and the medium was changed with RPMI containing 10% fetal bovine serum (Gibco, ThermoFisher), 100 units/mL penicillin/streptomycin (Gibco, ThermoFisher), 10 ng/mL human GM-CSF (PeproTech) and 10 ng/mL human IL-4 (PeproTech). In the case of tolDCs, 100 nM dexamethasone (Sigma-Aldrich) was also added to the medium.

For cell stimulation, LPS (10 ng/μL) and IFNγ (20 ng/μL) were added to cell culture at day 5, 24h before cell collection.

### CD8+ proliferation suppression assay

Allogenic CD8+ were isolated by negative selection using Dynabeads Untouched Human CD8 T Cells Kit (Invitrogen) and labeled with carboxyfluorescein succinimidyl ester (CFSE) CellTrace™ (Invitrogen), in accordance with the manufacturer’s instructions.

Purified CD8+ cells were seeded in 96-well plates (200,000 cells per well), and monocyte-derived cells (DCs or tolDCs) were added at different cell ratios (1:2, 1:3, 1:4, and 1:8). To stimulate CD8+ cells, 5 µL/mL of anti-CD3/CD28 Dynabeads (Invitrogen) were added to each well, except for the negative control.

For this experiment, cells were harvested in RPMI containing 10% fetal bovine serum (Gibco, ThermoFisher) and 100 units/mL penicillin/streptomycin (Gibco, ThermoFisher). Cells were cultured for 5 days, and the medium was changed on day 3.

### Quantification of cytokine production

Cell culture supernatants were collected after 6 days and diluted appropriately. Enzyme-linked immunosorbent assays (ELISA) were performed, following the manufacturer’s instructions: Human IL-10, Human IL-12p70, and Human TNFα from BioLegend, and Human IL-1β from ThermoFisher.

### Flow cytometry

To study cell-surface markers, cells were collected using Versene, a non-enzymatic dissociation buffer (ThermoFisher). Cells were resuspended in the staining buffer (PBS with 4% fetal bovine serum and 2 mM ethylenediaminetetraacetic acid (EDTA)). Cells were then incubated in ice with Fc block reagent (Miltenyi Biotec) for 10 minutes, and stained with the viability dye LIVE/DEAD™ Fixable Violet (ThermoFisher), following the manufacturer’s protocol.

Cells were then stained to study the proteins of interest, using the following antibodies: CD16 (APC) (#130-113-389, Miltenyi Biotec), CD14 (APC) (#130-110-520, Miltenyi Biotec), CD163 (FITC) (#33618, BioLegend), CD1a (PE) (#300106, BioLegend), CD80 (PE)(#H12208P, eBioScience), CD83 (APC) (#130-110-504, Miltenyi Biotec), CD86 (APC) (#130-113-569, Miltenyi Biotec), HLA-DR (PE) (#12-9956-42, eBioScience).

After staining, cells were fixed with PBS + 4% paraformaldehyde (Electron Microscopy Sciences) and analyzed within 2 days using a BD FACSCanto™ II Cell Analyzer (BD Biosciences). Data were analyzed with the FlowJo v10 software.

### Genomic DNA and total RNA extraction

Genomic DNA and total RNA were extracted using the Maxwell RSC Cultured Cells DNA kit (Promega) and the Maxwell RSC simplyRNA Cells kit (Promega), respectively, following the manufacturer’s instructions.

### Gene expression microarrays

RNA samples were processed in the Genomics Platform of the Vall d’Hebron Research Institute (Barcelona). After ensuring the RNA quality using Bioanalyzer, samples were hybridized in Clariom™ S microarrays (ThermoFisher).

Raw microarray data (CEL files) were analyzed using the oligo and limma packages of the Bioconductor project ^41^. First, raw data were normalized using the Robust Multichip Average algorithm (RMA), which is included as a function in the oligo package. We then performed an independent filtering step, removing probes with fewer than 3 samples over the 75% percentile of the negative control probes. After collapsing the remaining probes by gene, using the aggregate and mean functions, we built a limma linear model using the condition and donor information, with the formula ‘∼0+condition+donor’. The eBayes function in limma was then used in each pairwise comparison to obtain the FDR and logFC of each gene. Genes were considered to be differentially expressed when the FDR was less than 0.05 and the absolute logFC was greater than 0.5.

### Bisulfite pyrosequencing

500 ng of genomic DNA was converted using the EZ DNA Methylation Gold kit (Zymo Research). PCR was performed using the bisulfite-converted DNA as input and primers designed for each amplicon (Supplementary Table 5). These primers were designed using the PyroMark Assay Design 2.0 software (Qiagen). PCR amplicons were pyrosequenced using the PyroMark Q48 system and analyzed with PyroMark Q48 Autoprep software.

### Real-time quantitative polymerase chain reaction (RT-qPCR)

300ng of total RNA were reverse-transcribed to cDNA with Transcriptor First Strand cDNA Synthesis Kit (Roche) following manufacturer’s instructions. qRT-PCR was performed in technical triplicates, using LightCycler® 480 SYBR Green Mix (Roche), and 7.5ng of cDNA per reaction. The standard double-delta Ct method was used to determine the relative quantities of target genes, and values were normalized against the control genes RPL38 and HPRT1. Custom primers were designed to analyze genes of interest (Supplementary Table 5)

### DNA methylation profiling

500 ng of genomic DNA was converted using the EZ DNA Methylation Gold kit (Zymo Research). Infinium MethylationEPIC BeadChip (Illumina) arrays were used to analyze DNA methylation, following the manufacturer’s instructions. This platform allows around 850,000 methylation sites per sample to be interrogated at single-nucleotide resolution, covering 99% of the reference sequence (RefSeq) genes. Raw files (IDAT files) were provided for the Josep Carreras Research Institute Genomics Platform (Barcelona).

Quality control and analysis of EPIC arrays were performed using ShinyÉPICo ^42^, a graphical pipeline that uses minfi ^43^ for normalization, and limma ^41^ for differentially methylated positions analysis. CpH and SNP loci were removed and the Noob+Quantile normalization method was used. Donor information was used as a covariate, and Trend and Robust options were enabled for the eBayes moderated t-test analysis. CpGs were considered differentially methylated when the absolute differential of methylation was greater than 20% and the FDR was less than 0.05.

### Immunofluorescence

Cells were fixed with PBS + 4% paraformaldehyde for 20 min and permeabilized with PBS + Triton X-100 0.5% for 10 min. Coverslips were washed twice with PBS and blocked with PBS + 4% BSA for 1 h. Anti-MAFB antibody HPA005653 (Sigma-Aldrich) was diluted 1:100 in PBS and incubated overnight with the samples in a humidity chamber. Then, cells were incubated with anti-rabbit Alexa Fluor 647 (ThermoFisher) in blocking solution (PBS + BSA 4% + 0.025% Tween 20) for 1 h. After four washes with PBS, cells were stained with Alexa Fluor™ 488 Phalloidin (ThermoFisher) 1/200 and DAPI 2 µg/mL. Vectashield (Vector Laboratories) was used for the final sample preparation in slides. Images were obtained with a Leica TCS-SL confocal microscope.

### Western blotting

Cytoplasmic and nuclear protein fractions were obtained using hypotonic lysis buffer (Buffer A; 10 mM Tris pH 7.9, 1.5 mM MgCl2, 10 mM KCl supplemented with protease inhibitor cocktail (Complete, Roche) and phosphatase inhibitor cocktail (PhosSTOP, Roche) to lyse the plasma membrane. Cells were visualized under the microscope to ensure correct cell lysis. Nuclear pellets were resuspended in Laemmli 1X loading buffer. For whole-cell protein extract, cell pellets were directly resuspended in Laemmli 1X loading buffer.

Proteins were separated by SDS-PAGE electrophoresis. Immunoblotting was performed on polyvinylidene difluoride (PVDF) membranes following standard procedures. Membranes were blocked with 5% Difco™ Skim Milk (BD Biosciences) and blotted with primary antibodies. After overnight incubation, membranes were washed three times for 10 min with TBS-T (50 mM Tris, 150 mM NaCl, 0.1% Tween-20) and incubated for 1 h with HRP-conjugated mouse or rabbit secondary antibody solutions (Thermo Fisher) diluted in 5% milk (diluted 1/10000). Finally, proteins were detected by chemiluminescence using WesternBright™ ECL (Advansta). The following antibodies were used: Anti-MAFB (HPA005653, Sigma-Aldrich), Anti-GR (C15200010-50, Diagenode), Anti-GAPDH (2275-PC-100, Trevigen), Anti-Lamin B1 (ab229025, Abcam), Anti-TET2 (C15200179, Diagenode).

### Co-immunoprecipitation (Co-IP)

Co-IP assays were performed using tolDCs differentiated from CD14+ monocytes for 24 h. Cell extracts were prepared in lysis buffer [50 mM Tris–HCl, pH 7.5, 1 mM EDTA, 150 mM NaCl, 1% Triton-X-100, protease inhibitor cocktail (cOmplete™, Merck)] with corresponding units of Benzonase (Sigma) and incubated at 4°C for 4 h. 100 μL of supernatant was saved as input and diluted with 2× Laemmli sample buffer (5x SDS, 20% glycerol, 1M Tris–HCl (pH 8.1)). Supernatants were first precleared with PureProteome™ Protein A/G agarose suspension (Merck Millipore) for 1 h. The lysate was then incubated overnight at 4°C with respective crosslinked primary antibodies. Cross-linking was performed in 20 mM dimethyl pimelimidate (DMP) (Pierce, ThermoFisher Scientific, MA, USA) dissolved in 0.2 M sodium borate (pH 9.0). Subsequently, the beads were quenched with 0.2M of ethanolamine (pH 8.0) and resuspended at 4°C in PBS until use. Beads were then washed three times with lysis buffer at 4°C. Samples were eluted by acidification using a buffer containing 0.2 M glycine (pH 2.3) and diluted with 2× Laemmli. Samples and inputs were denatured at 95°C in the presence of 1% β-mercaptoethanol. Anti-MAFB HPA005653 (Sigma-Aldrich), Anti-GR C15200010-50, and Anti-TET2 antibody ab124297 (Abcam) were used for Co-IP.

### Transfection of primary human monocytes

We used the #si19279 MAFB Silencer Select siRNA (Agilent, ThermoFisher) to perform knockdown experiments in MOs, using the Silencer Select Negative Control #1 (Agilent, ThermoFisher) as control. CD14+ MOs were cultured in 12-well plates (1 million cells/well) and transfected with siRNAs (100 nM) using Lipofectamine 3000 Reagent (1.5 µL/well), following the manufacturer’s protocol. After 6 h, the medium was changed and DCs/tolDCs were cultured as described in “CD14+ monocytes purification and culture”. To test transfection efficiency, siGLO Green Transfection Indicator (Horizon) was used, at the same concentration, and checked by flow cytometry 24 h after transfection.

### RNA-seq

RNA-seq libraries of transfected DC/tolDCs were generated and sequenced by BGI Genomics (Hong Kong), in 100-bp paired-end, with the DNBseq platform. More than 30 million reads were obtained for each sample. Fastq files were aligned to the hg38 transcriptome using HISAT2^44^ with standard options. Reads mapped in proper pair and primary alignments were selected with SAMtools ^45^. Reads were assigned to genes with featureCounts ^46^.

Differentially expressed genes were detected with DESeq2 ^47^. The donor was used as a covariate in the model. The Ashr shrinkage algorithm was applied and only protein-coding genes with an absolute logFC greater than 0.5 and an FDR less than 0.05 were selected as differentially expressed. For representation purposes, Variance Stabilizing Transformation (VST) values and normalized counts provided by DESeq2 were used.

### Chromatin immunoprecipitation

After 4 or 24 h of cell culture, DCs and tolDCs were fixed with Pierce™ fresh methanol-free formaldehyde (ThermoFisher) for 15 min and prepared for sonication with the truChIP Chromatin Shearing Kit (Covaris), following the manufacturer’s instructions. Chromatin was sonicated for 18 min with the Covaris M220 in 1mL milliTubes (Covaris). Size distribution of the sonicated chromatin was checked by electrophoresis to ensure appropriate sonication, with a size of around 200 bp. Magna Beads Protein A+G (Millipore) was blocked with PBS + BSA (5 mg/mL) for 1 h. Chromatin was precleared with 25 µl of beads for 1.5 h and 10 µg of chromatin were incubated overnight with each antibody: 10 µl Anti-MAFB antibody HPA005653 (Sigma-Aldrich) and 5 µl Anti-GR antibody C15200010-50 (Diagenode), in a buffer with 1% Triton X-100 and 150 mM NaCl.

Three washes were performed with the Low Salt Wash Buffer (0.1% SDS, 1%Triton X-100, 2 mM EDTA, pH 8.0, 20 mM Tris-HCl pH 8.0, 150 mM NaCl), the High Salt Wash Buffer (0.1%SDS, 1% Triton X-100, 2 mM EDTA, pH 8.0, 20 mM Tris-HCl pH 8, 500 mM NaCl), and the LiCl Wash Buffer (0.25 M LiCl, 1% Nonidet P-40, 1% Deoxycholate, 1 mM EDTA pH 8, 10 mM Tris-HCl), followed by a final wash with TE buffer (pH 8.0, 10 mM Tris-HCl, 1 mM EDTA). Chromatin was eluted for 45 min at 65°C with 100 µl of elution buffer (10 mM Tris-Cl, 1 mM EDTA, 1% SDS) and decrosslinked by adding 5 µl 5M NaCl and 5 µl 1M NaHCO_3_ (2 h at 65°C). Next, 1 µl of 10 mg/mL proteinase K (Invitrogen) was added and the samples were incubated at 37°C for 1 h.

For DNA purification, iPure kit v2 (Diagenode) was used, following the manufacturer’s instructions.

### Chromatin immunoprecipitation sequencing (ChIP-seq) analysis

ChIP-seq inputs and immunoprecipitated DNA samples were used to generate ChIP-seq libraries in the Centre for Genomic Regulation (Barcelona), using the TruSeq ChIP Library Preparation Kit (Illumina). Quality control of the libraries was performed using 2100 Bioanalyzer (Agilent). Libraries were sequenced with an Illumina HiSeq 2500, in 50-bp single-end, yielding between 25 and 40 million reads per sample.

Potential adapter contamination was trimmed from the raw reads using Cutadapt (https://doi.org/10.14806/ej.17.1.200). Reads were aligned to the GRCh38 human genome assembly using the Burrows-Wheeler Alignment (BWA)-MEM algorithm ^48^. Aligned reads were filtered by MAPQ, removing alignments with MAPQ < 30, using SAMtools ^45^. Aligned reads overlapping with the ENCODE blacklist were also removed ^49^.

Quality control of ChIP-seq data was performed using the SPP package^50^. The relative strand cross-correlation coefficient (RSC) was greater than or equal to 2 in all the immunoprecipitated samples.

Bigwig files were generated for visualization, using the bamCoverage function in the Deeptools package ^51^, with the bins per million mapped reads (BPM) method and ‘--binSize 20 --extendReads 150 --smoothLength 60 --centerReads’ options. Wiggletools was used to aggregate the bigwig duplicates ^52^.

MACS2 software with ‘--nomodel --extsize 200’ options was used for peak calling ^53^. Duplicates were aggregated using the MSPC algorithm ^54^ with the options ‘-r Biological -w 1E-6 -s 1E-12’. The resulting consensus peaks were used for the downstream analysis.

### Data analysis and representation

Statistical analyses were performed in R 4.0. Gene expression and DNA methylation heatmaps were created with the heatmap.2 function of the gplots package. The findMotifsGenome.pl function of HOMER (Hypergeometric Optimization of Motif EnRichment) was used to analyze known and *de novo* motif enrichment. For ChIP-seq peaks, the parameter ‘-size 50’ was used, whereas the parameters ‘-size 200 -cpg) were used for methylation data. All EPIC array CpG coordinates were also used as background for the methylation data. GREAT software was used to calculate CpG-associated genes and gene ontology (GO) enrichment ^55^. Gene set enrichment analysis (GSEA) and GO enrichment of gene expression data were performed using the clusterProfiler package ^56^. ChIP-seq peaks files of histone marks from MO and DCs were downloaded from the BLUEprint webpage (http://dcc.blueprint-epigenome.eu). Consensus peaks of the different replicates were obtained with the MSPC algorithm, using the options ‘-r Biological -w 1E-4 -s 1E-8 -c 3’.

The chromatin state learning model for CD14+ monocytes was downloaded from the Roadmap Epigenomics Project webpage, and chromatin state enrichments were calculated using Fisher’s exact test.

ChIP-seq overlaps and Venn diagrams were generated with the ChIPpeakAnno package ^57^. Genomic track plots were created with pyGenomeTracks ^58^. Methylated CpG set enrichment analysis (mCSEA) ^59^ was used to calculate CpG-set-specific DNA methylation modifications. Public GR ChIP-seq datasets were extracted from the Remap2020 database ^60^, excluding samples without glucocorticoid treatment. Peak intersects were calculated using bedtools ^61^. Public peak-callings of histone marks were extracted from the Blueprint database^22^ and replicates were aggregated using the MSPC algorithm. Public DNAse-seq and ATAC-seq bigwigs were aggregated using wiggletools.

### Student’s paired t test

Statistical analyses involved Student’s paired-samples t tests, with which the means of matched pairs of groups were compared, except where indicated otherwise. The levels of significance were: ns, p > 0.05; *, p < 0.05; **, p < 0.01; ***, p < 0.001.

### Gene expression public data processing

To compare gene expression with public data, expression array matrices were downloaded from the GEO database (GSE40484, GSE117946 and GSE99056). Our expression dataset and public expression data were combined by gene symbol and batch-corrected using ComBat ^62^. The 1000 most variable genes were used to plot the data in a T-distributed Stochastic Neighbor Embedding (t-SNE) representation, using the Rtsne package (https://www.jmlr.org/papers/v9/vandermaaten08a.html).

### Synovial tissue extraction and processing

Synovial biopsy and tissue analysis was approved by the Kantonal Ethic Commission Zurich, Switzerland (refs: 2019-00115 and 2016-02014). All patients signed informed consent forms. Patient’s characteristics were as follows:

|  | Gender | Site | Diagnosis | Rheumatoid factor | Synovitis score | Histology | Therapy |
| --- | --- | --- | --- | --- | --- | --- | --- |
| 28 | f | MCP | RA | negative | 5 | lympho-myeloid | Glucocorticoids oral 10mg/d |
| 50 | f | wrist | RA | negative | 8 | lympho-myeloid | none |

One part of the synovial biopsies was embedded in paraffin and histologically analyzed as previously described ^63, 64^. For scRNA-seq synovial biopsies were mechanically minced and enzymatically digested with 100 ug/ml Liberase TL (Roche) and 100 ug/ml DNAseI (Roche) in pre-warmed RPMI 1640 containing 2 mM glutamine and 25 mM HEPESGibco for 30 min at 37°C. The digestion was stopped by adding 20 v/v % fetal calf serum (FCS) and the dissociated tissue was filtered through a 40 um cell strainer. Red blood cells were lysed with RBC lysis buffer (Milteny Biotec). Cell viability was analyzed with a LUNA-FL dual fluorescence cell counter (88% and 90%).

### Single-cell RNA-seq processing and analysis

Dissociated cells were processed with the Chromium Single Cell 3ʹ GEM, Library & Gel Bead Kit v3 or v3.1 according to the manufacturer’s protocol (10x Genomics). Libraries were sequenced on the NovaSeq 6000 (Illumina). Raw sequencing data was processed with cell ranger (v6.0.0, 10x Genomics) using mkfastq and count with default settings and the provided human reference (GRCh38-2020-A).

|  | Targeted cell number | Estimated number of cells | Reads/cell |
| --- | --- | --- | --- |
| 28 | 10000 | 4389 | 44396 |
| 50 | 6000 | 5760 | 53176 |

We used Seurat (4.0.3) to analyze scRNA-seq data. We first read and merge data from patients, and we filtered cells with counts lower than 200 or higher than 30000 and with RNA features lower than 100 or higher than 5000. We also filtered cells with mitochondrial or *r*ibosomal features higher than 25%. We then integrated samples before clustering, using the standard Seurat pipeline. First, features were normalized and variable features were found for each sample, independently. Next, functions FindIntegrationAnchors and IntegrateData were used to integrate the data. The functions ScaleData (using RNA counts and mitochondrial percentage as variables to regress), RunPCA, FindNeigbors, FindClusters and RunUMAP were used sequentially to perform the UMAP representation, with 30 dimensions. After selecting only monocyte-derived clusters, scaling and representation was performed again, using the same functions, with resolution 0.15 and 7 dimensions.

## Data access

DNA methylation, expression and ChIP-seq data for this publication have been deposited in the NCBI Gene Expression Omnibus and are accessible through GEO SuperSeries accession number GSE180542

## Acknowledgements

We are very grateful to Prof. Angel Corbí and his team for useful discussions and feedback.

## Author contributions

O.M.-P. and E.B. conceived and designed the study; O.M.-P., L.C., C.D.-F., T.L., M.H., S.E., M.F.-B. and G.B. performed experiments; O.M.-P. performed the bioinformatic analyses; R.M., M.H., S.E., M.F.-B. and C.O. generated and analyzed the single cell datasets; O.M.-P., R.M., C.O. and E.B. analyzed results; O.M.-P. and E.B. wrote the manuscript; all authors participated in discussions and in interpreting the results.

